# CRAVAT 4: Cancer-Related Analysis of Variants Toolkit

**DOI:** 10.1101/162859

**Authors:** David L. Masica, Christopher Douville, Collin Tokheim, Rohit Bhattacharya, RyangGuk Kim, Kyle Moad, Michael C. Ryan, Rachel Karchin

**Affiliations:** Department of Biomedical Engineering, The Johns Hopkins University, Baltimore, MD; The Institute for Computational Medicine, The Johns Hopkins University, Baltimore, MD; Department of Computer Science, The Johns Hopkins University, Baltimore, MD; In Silico Solutions, Falls Church, VA; Department of Oncology, The Johns Hopkins University School of Medicine, Baltimore, MD

**Keywords:** New software for data analysis, Suppressor genes, Oncogenes, CRAVAT, MuPIT, VEST

## Abstract

Cancer sequencing studies are increasingly comprehensive and well-powered, returning long lists of somatic mutations that can be difficult to sort and interpret. Diligent analysis and quality control can require multiple computational tools of distinct utility and producing disparate output, creating additional challenges for the investigator. The Cancer-Related Analysis of Variants Toolkit (CRAVAT) is an evolving suite of informatics tools for mutation interpretation that includes mutation projecting and quality control, impact prediction and extensive annotation, gene- and mutation-level interpretation including joint prioritization of all nonsilent consequence types, and structural and mechanistic visualization. Results from CRAVAT submissions are explored in an interactive, user-friendly web-environment with dynamic filtering and sorting designed to highlight the most informative mutation, even in the context of very large studies. CRAVAT can be run on a public web-portal, in the cloud, or downloaded for local use, and is easily integrated with other methods for cancer omics analysis.

**Conflict of interest:** All authors declare no potential conflict of interest.

## Background

An investigator’s work is far from over when results are returned from the sequencing center. Depending on the service, genetic mutation calls can require additional projecting to include all relevant RNA transcripts or correct protein sequences. Assignment of consequence type or sequence ontology (e.g., missense, splice, or indel) might be incomplete. Once projecting is complete and sequence ontology assigned, the task of interpreting mutation impact remains. This interpretation can entail annotation from multiple sources, employing one or more bioinformatics classifiers, testing for functional or pathway enrichment, and referencing each mutation against databases of known cancer drivers or common polymorphisms. Making full use of these diverse resources can be a daunting task. The individual utilities often assess a limited number of consequence types, require different formatting of input data, and return disparate output not amenable to direct comparison. Once the investigator has managed to assemble a suitable *ad hoc* pipeline to process and interpret each mutation, the task of effectively sorting and filtering large lists of mutations may be difficult.

CRAVAT^1^ is designed to streamline the many steps outlined above, ultimately returning mutation interpretations in a dynamic interactive web environment for sorting, visualizing, and inferring mechanism. CRAVAT is suitable for large studies (e.g., full-exome and large cohorts) and small studies (e.g., gene panel or single patient), performs all projecting and assigns sequence ontology, predicts mutation impact using multiple bioinformatics classifiers normalized to provide comparable output, allows for joint prioritization of all nonsilent mutation types, organizes annotation from many sources on graphical displays of protein sequence and 3D structure, and facilitates dynamic filtering and sorting of results so that the most important mutations can be quickly retrieved from large submissions. In addition to running on our webserver, CRAVAT can be dockerized for cloud computing or downloaded for local use, and is easily integrated with other software packages. The following describes the details of the CRAVAT interactive results explorer; machine-readable text and Excel spreadsheets are also provided with each job submission.

## CRAVAT 4.x

The CRAVAT webserver (cravat.us) prompts the user to submit sequencing data, select impact prediction methods, and provide an email address for communicating job status. Once sequencing data is submitted, CRAVAT performs quality control and projects all mutations from genome to transcript—projecting each mutation to all relevant transcripts—and then to protein sequence; additionally, mutations are projected to available protein structures and homology models. Next, all consequence types (sequence ontologies) are assigned, including missense, stop gains and losses, in-frame and frame-shifting insertions and deletions (indels), splice, and silent mutations.

### Mutation impact prediction

CRAVAT employs two approaches for predicting mutation impact, namely CHASM (Cancer-specific High-throughput Annotation of Somatic Mutations)^2^ and VEST (Variant Effect Scoring Tool).^3^, ^4^ CHASM uses a Random Forest classifier that is trained from a positive class of cancer driver mutations from COSMIC^5^ and a putative set of passenger mutations; thus, CHASM classifies mutations as cancer *drivers* or *passengers*. VEST uses a Random Forest classifier that is trained from a positive class of disease-associated germline mutations from the Human Gene Mutation Database (HGMD)^6^ and a negative class of common mutations from the exome sequencing project^7^; thus, VEST classifies mutations as *pathogenic* or *benign*. CHASM can score missense mutations. VEST scores all nonsilent consequence types, and facilitates joint prioritization (i.e., scores are directly comparable among all mutations, regardless of sequence ontology). Both CHASM and VEST provide composite gene-level p-values and false discovery rates (FDRs), as well as a mutation-specific p-values and FDRs.

### Results Summary

Figure 1A shows selected displays from the CRAVAT *Summary* tab. This summary includes aggregate results for all mutations from a single submission, at the patient and cohort level. Distribution of submitted mutations by gene function and sequence ontology are displayed as pie charts (see Figure 1A); a pie chart showing distribution by Cancer Genome Landscape definitions^8^ is also provided. Chromosome-specific distribution of nonsilent, missense, and inactivating mutations are displayed on a circos plot (see Figure 1A). The results summary also shows the most frequently mutated genes (corrected for gene length) as a bar chart, the top-10 most significant genes as determined by VEST or CHASM composite p-values, and sample-level consequence-type distributions displayed as a series of stacked histograms (see Figure 1A). Gene-set enrichment—of collections of genes with significant composite VEST or CHASM p-values—is embedded as NDEx networks^9^; here, the user can toggle through each network detected for their submission and view significant genes in the context of larger biological networks (see Figure 1A).

**Figure 1.**
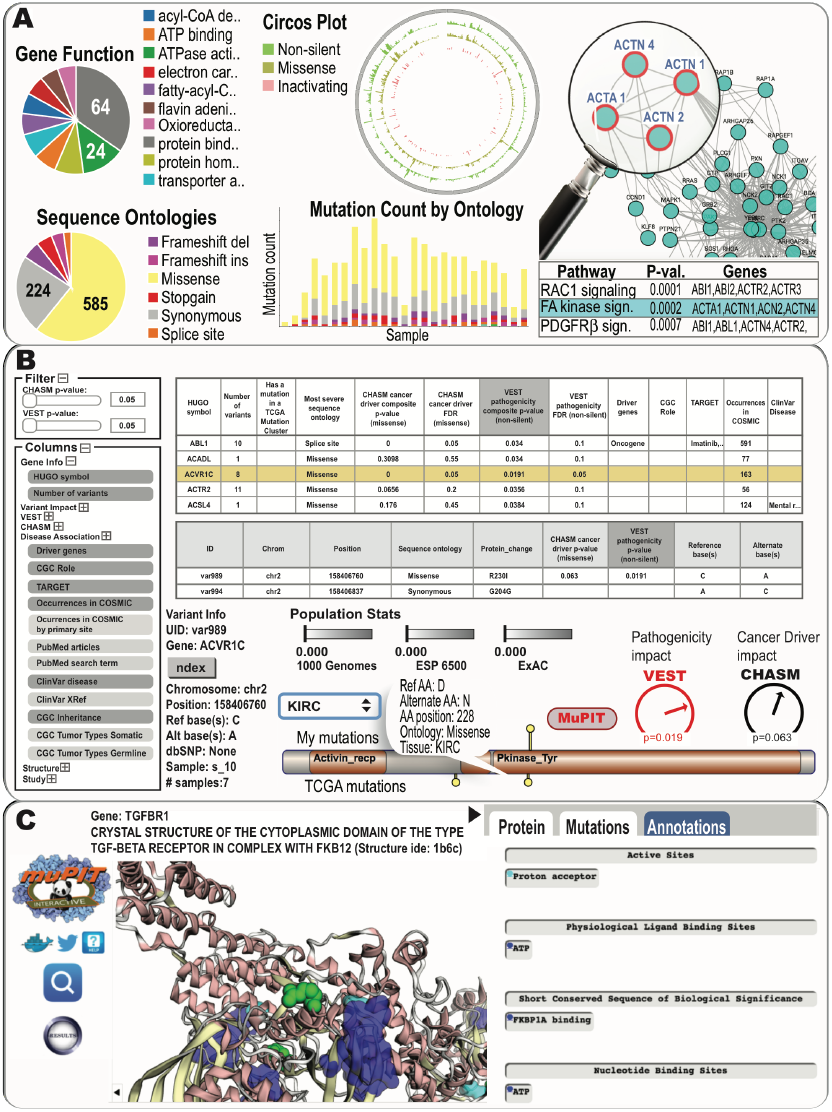
Selected widgets and displays from the CRAVAT Summary tab (A), the Gene and Variant tabs (B), and from MuPIT (C).

### Gene- and Mutation-level analysis

Figure 1B shows selected displays from the CRAVAT *Gene* and *Variant* tabs. The *Gene* tab displays an interactive spreadsheet including every gene that harbored a mutation from the submission (truncated example shown at top of Figure 1B). This table includes gene-specific mutation frequency, the most severe consequence type among submitted mutations for each gene, presence of statistically significant mutation clustering in the related protein (inferred from TCGA data, see below), composite VEST and CHASM p-values and FDRs, classification as a tumor suppressor or oncogene if applicable, associated drugs, COSMIC occurrences, ClinVar disease status, and Gene Ontology (GO) terms. The table can be sorted by any category and filtered by CHASM and VEST significance (see filtering widget, Figure 1B); any instantiation of this table can be exported in Excel format. Clicking any row (i.e., gene) in this table retrieves a drill-down table and a lollipop chart. The drill-down table displays information for each mutation from the selected gene, including chromosome position, sequence ontology, protein change, mutation-specific VEST and CHASM p-values, and the reference and alternative DNA base(s). The lollipop diagram plots all submitted mutations for the selected gene onto the protein sequence, with domain labels and mutation-specific information accessible through the tooltip (mouse-over); the user can optionally choose to plot tissue-specific mutations from TCGA data.

The CRAVAT interactive *Variant* tab serves information that will be familiar to users after exploring the *Gene* tab, but here the user can drill down into the details even further (see for instance, the second spreadsheet in Figure 1B). This tab includes p-values and FDRs for each mutation in the context of all relevant transcripts. Frequencies from population databases (1000 Genomes^10^, ESP6500^7^, and ExAC^11^) are graphically displayed, and can be further partitioned by ethnicity. Figure 1B shows an example lollipop diagram, where the submitted R230I missense substitution (top side of lollipop diagram) is compared with TCGA kidney cancer data (bottom of lollipop diagram), revealing the nearby D228N substitution in the *ACVR1C* gene.

### MuPIT Interactive

Figure 1C shows selected displays from MuPIT (Mutation Position Imaging Toolbox) Interactive.^12^ MuPIT projects mutation positions to annotated, interactive 3D protein structures, and links directly from the gene and mutation entries in the above-described widgets and tables. MuPIT achieves high coverage by including experimental structures from the protein data bank and homology models filtered using standard measures for model quality. MuPIT was designed for investigators without expertise in protein structure/function, and can be used to develop mechanistic hypotheses regarding mutations of interest.

For each structure, the MuPIT *Protein* tab allows manipulation of graphical parameters such as color, outline, opacity, and style (cartoon, stick, space fill, etc.), as well as options for displaying specific domains, chains, and ligands; additionally, publication-ready images can be exported from this tab. The *Mutations* tab allows the user to toggle through submitted mutations alongside statistically significant 3D mutation clusters from TCGA data; statistically significant mutation clusters are precomputed for each of 31 TCGA cancer subtypes using the HotMAPS algorithm.^13^ In Figure 1C, MuPIT is located at the *Annotations* tab, and user-submitted mutations (shown in green) are spatially proximal to known TGFBR1 binding and active sites (cyan and blue), suggesting a potential mechanistic role for these mutations.

## CRAVAT 5

The next update of CRAVAT is scheduled for release in 2017, and will include annotations for noncoding mutations, detection of statistically significant sequence hotspots for mutations, updated VEST and CHASM classifiers, inclusion of top-performing impact prediction algorithms from other groups, migration to hg38, and integration with the Broad Integrated Genomics Viewer (IGV). The corresponding MuPIT update will include pathogenic germline mutations, and rare and common germline mutation from healthy populations, for optional projecting and display onto protein structure.

### In context with other omics analysis

CRAVAT is developed for easy integration with other software, including programmatic interfaces for web-services. This flexibility affords the combination of CRAVAT’s mutation interpretation capabilities with software considering other omics datatypes (e.g., copy-number alterations, methylation, expression). For instance, other methods currently integrating CRAVAT include: Trinity, which assembles Illumina RNA-seq data^14^; UCSC Xena, which can combine many omics and clinical datatypes^15^; the Network Data Exchange (NDEx), an online commons for biological network data.^9^ CRAVAT can also be used with Galaxy tools.

